# Improved Sensitivity in Low-Input Proteomics using Micro-Pillar Array-based Chromatography

**DOI:** 10.1101/678995

**Authors:** Johannes Stadlmann, Otto Hudecz, Gabriela Krššáková, Gert Van Raemdonck, Jeff Op De Beeck, Gert Desmet, Josef M. Penninger, Paul Jacobs, Karl Mechtler

## Abstract

Capitalizing on the massive increase in sample concentrations which are produced by extremely low elution volumes, nano-LC-ESI-MS/MS is currently one of the most sensitive analytical technologies for the comprehensive characterization of complex protein samples.

However, despite tremendous technological improvements made in the production and the packing of monodisperse spherical particles for nano-flow HPLC, current state-of-the-art systems still suffer from limits in operation at the maximum potential of the technology.

With the recent introduction of the µPAC system, which provides perfectly ordered micro-pillar array based chromatographic support materials, completely new chromatographic concepts for optimization towards the needs of ultra-sensitive proteomics become available.

Here we report on a series of benchmarking experiments comparing the performance of a commercially available 50 cm micro-pillar array column to a widely used nano-flow HPLC column for the proteomics analysis of 10 ng tryptic HeLa cell digest.

Comparative analysis of LC-MS/MS-data corroborated that micro-pillar array cartridges provide outstanding chromatographic performance, excellent retention time stability, increase sensitivity in the analysis of low-input proteomics samples, and thus repeatedly yielded almost twice as many unique peptide and unique protein group identifications when compared to conventional nano-flow HPLC columns.

## TECHNICAL BRIEF

The field of proteomics aims at the qualitative and quantitative description of all proteins contained in complex biological samples. Currently, the most sensitive proteomics platforms are almost exclusively based on the combination of two key analytical methods: nano-flow High Performance Liquid Chromatography (nano-HPLC) and tandem mass-spectrometry (MS/MS), hyphenated by ElectroSpray-Ionization (ESI).

However, despite the tremendous improvements in ultra-sensitive nano-HPLC-ESI-MS/MS-based proteomics workflows and instrumentation, which capitalize on the massive increase in sample concentrations produced by extremely low elution volumes, the comprehensive characterization of e.g. single mammalian cells still challenges the sensitivity of currently available technologies.

Next to the development of dedicated low-input sample preparation methods [1-4], which aim at reducing sample losses prior to analysis, changes to the chromatographic support material have recently been identified key to the sensitive profiling of low-input proteomics samples by nano-HPLC-ESI-MS/MS [5,6].

While nonporous particles have been demonstrated to hold the great potential of high chromatographic separation power and to further reduce on-column losses [5], their expedient integration into standard proteomics nano-flow HPLC systems has long suffered from their intrinsically low loading capacity [6]. However, especially in the context of low-input proteomics, which aims at analyzing protein amounts in the nano-to pico-gram-range, the capacity of these novel reversed-phase (RP) HPLC support materials is not a limiting factor. More importantly, the potential gain in sensitivity due to improved peak width and peak capacities provided by these new types of chromatographic columns, makes them extremely attractive for ultrasensitive proteomics applications.

With the recent commercialization of perfectly ordered micropillar array-based nano-HPLC cartridges (µPAC, PharmaFluidics), we wished to explore potential benefits of this technology to the ultra-sensitive analysis of low-input proteomics samples. Here we report on a series of benchmarking experiments comparing the performance of a 50 cm micro-pillar array nano-HPLC cartridge to a state-of-the-art, particle-based nano-HPLC column, in the analysis of 10 ng tryptic HeLa cell protein digest.

For these experiments, we installed the respective pre- and analytical columns (i.e. µPAC RP18, 50 cm, PharmaFluidics; PepMap C18, 3 μm, 75 µm × 50 cm, Thermo) in identical LC-ESI-MS/MS setups, all comprising an Ultimate 3000 RSLCnano LC system (Dionex – Thermo) operated at 50 °C, coupled to the exact same Q Exactive HF-X mass-spectrometer (Thermo). The samples (10 ng/µL HeLa digest, Pierce; in 0.1% formic acid) were injected using a 1µL sample-loop at full loop injection, were trapped on a pre-column and then separated by developing two-step linear gradients of increasing length, at a fixed flow-rate of 250 nL/min: from 2% to 20% acetonitrile in 0.1% formic acid in 45, 90 and 135 min., followed by 20% to 32% acetonitrile in 0.1% formic acid within 15, 30 and 45 min (i.e. 60, 120 or 180 min total gradient time), respectively. All three gradient programs were completed by a final gradient step from 32 to 78% acetonitrile in 0.1% formic acid, within 5 min.

The mass-spectrometer was operated in positive mode and set to the following acquisition parameters: MS1 resolution = 60000, MS1 AGC-target = 1E6, MS1 maximum inject time = 60 ms, MS1 scan range = 350-1500 m/z, MS2 resolution = 15000, 45000 or 60000, MS2 AGC-target = 2E5, maximum inject time = 105 ms, TopN = 10, isolation window = 0.7 m/z, MS2 scan range = “dynamic first mass”, normalized collision energy = 28, minimum AGC target =1E4, intensity threshold 9.5e4, pre-cursor charge states = 2-6, peptide match = preferred, exclude isotopes = ON, dynamic exclusion = 45 s, “if idle…” = do not pick others. All experiments were performed in technical triplicates.

Subsequently, all LC-MS/MS raw-data were processed and identified using Proteome Discoverer (version 2.3.0.523, Thermo Scientific). For this, MS/MS spectra were extracted from the raw-files and searched against the Swissprot protein database, restricting taxonomy to Homo sapiens and including common contaminant protein sequences (20341 sequences; 11361548 residues) using MSAmanda (Engine version v2.0.0.12368) [7]. The search engine parameters were set as follows: peptide mass tolerance= ±7 ppm, fragment mass tolerance= 15ppm, cleavage specificity= trypsin, missed cleavage sites= 2, fixed modifications= carbamidomethylation of cysteine, variable modifications= oxidation of methionine. Results of the MS/MS search engine were filtered to 1 % FDR on protein and peptide level using the Elutator algorithm [8], implemented as node to Proteome Discoverer 2.3. Identified peptide features were extracted from the raw-files and quantified, using the in-house-developed Proteome Discover-node apQuant [9].

Comparative analysis of the MS/MS data (Fig.1. A.,B.) highlighted that µPAC cartridges repeatedly yielded almost twice as many unique peptide identifications (e.g. 15629, at 120-min gradient length) and unique protein groups (e.g. 2743, at 120-min gradient length), when compared to the PepMap C18 system (e.g. 7364 unique peptides and 1500 unique protein groups, at a gradient length of 120 min). Of note, enabling the 2^nd^ search-option available in MS Amanda [7] yielded approx. 30% more peptide spectrum matches and approx. 20% more peptide identifications, irrespective of the chromatographic system used (data not shown). An overlay of the identifications showed excellent agreement with respect to a core-set of peptides (Fig.1. C.) and protein groups (Fig.1. D.), which were identified with both nano-HPLC setups. Remarkably, however, although >90% of all peptide and protein group identified with the PepMap C18 system were also found in the µPAC set-up, the latter allowed for the additional identification of 80% more peptides and 90% more protein groups (Fig.1. C., D.). This important increase in peptide and protein group identifications specifically provided by the µPAC system, were primarily due to almost twice as many MS/MS spectra triggered in the respective experiments (e.g. 26358 and 14027, at 120-min gradient length, for µPAC and PepMap, respectively), corroborating increased sensitivity. Of note, the maximum TopN number of 10 precursor ion-masses to be scheduled for MS/MS analysis was hardly reached in the µPAC experiments (e.g. in 10 duty-cycles, at 120-min gradient length), and never exceeded more than 6 precursor ion-masses to be scheduled for MS/MS in the PepMap experiments at 120-min gradient length.

**Figure 1.**
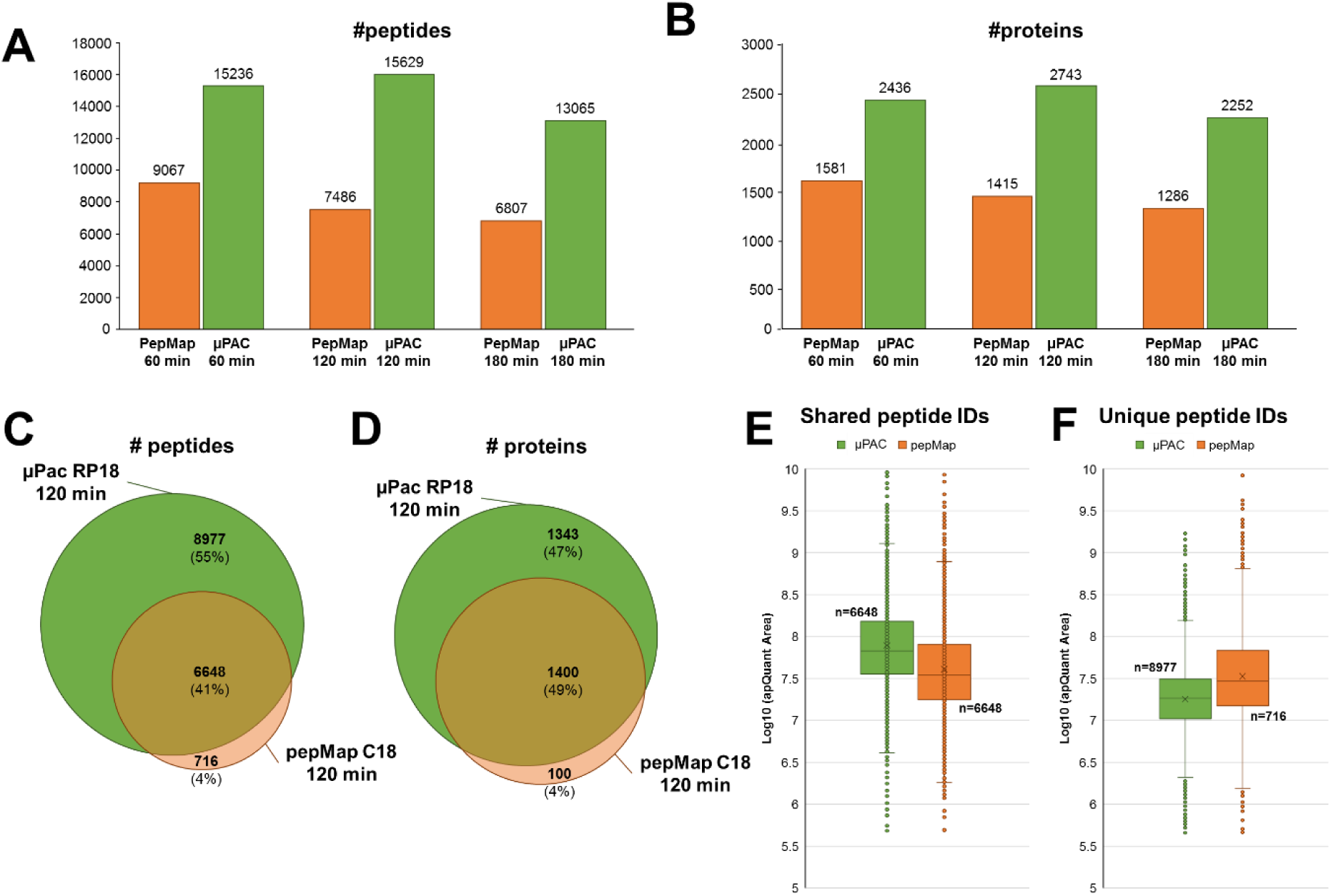
Comparative analysis of the LC-MS/MS data generated from 10 ng HeLa digest, using the µPAC RP18 or the PepMap C18 set-up. **A)** Number of unique peptide and **B)** unique protein groups identified, at different gradient lengths. All experiments were performed in technical triplicates. Overlay of repeatedly identified **C)** peptide sequences and **D)** protein groups. Comparative “Box-and-whiskers” plots of precursor-ion specific chromatographic peak areas of **E)** peptides identified in both nano-HPLC set-ups and **F)** peptides which were exclusively identified in either of the two nano-HPLC set-ups at 120 min. gradient length. All data processing, chromatographic peak detection and peak area calculations were performed using Proteome Discoverer 2.3, MS Amanda and apQuant.

Further investigating the substantial µPAC-specific gain in peptide and protein identifications, we extracted peptide precursor-ion specific chromatographs and calculated the respective chromatographic peak areas, using apQuant. Comparison of the data highlighted that peptide precursor-ion intensities were on average 2-fold higher when using the µPAC cartridges (Fig.1. E). By contrast, the µPAC-only identifications (i.e. 8977 peptides and 1343 proteins; Fig1. C.), predominantly derived from low abundant peptide precursor-ions (Fig.1. F.). The sparse PepMap-only identified peptides, however, were detected within a precursor-ion intensity range which was similar to common peptide identifications.

Next, we analyzed chromatographic performance parameters of the two systems. For this, we automatically extracted peptide-specific retention times at the peak apex, determined full-width at half-maximum (FWHM) and calculated peak asymmetry parameters (i.e. skewness), using apQuant. The comparison of peptide-specific peak widths (i.e. FWHM) revealed broader peaks on the PepMap columns (i.e. mean = 12.9 sec. and 10.2 sec, for a 120 min. gradient on PepMap and µPAC, respectively; Fig. 2. A.) and suggested a 25% increase in peak capacity for the µPAC system (i.e. estimated peak capacity of 560 and 705, at 120 min gradient length, for PepMap and µPAC, respectively). More importantly, however, while 90% of all peptides identified with µPAC cartridges had a FWHM of < 12 sec., the same proportion of peptides were detected with a FWHM of < 16 sec on the PepMap C18 system. (Fig. 2. B.). Additionally, while both nano-LC set-ups performed similarly at 60 min. gradient length (Fig.2. C.), the µPAC system provided superior peak-width at extended gradient times (i.e. 180 min.; Fig.2. D.). Chromatographic peak symmetry, as determined by calculating peptide-specific peak skewness, was found similar in both set-ups (Fig. 2. E.).

**Figure 2.**
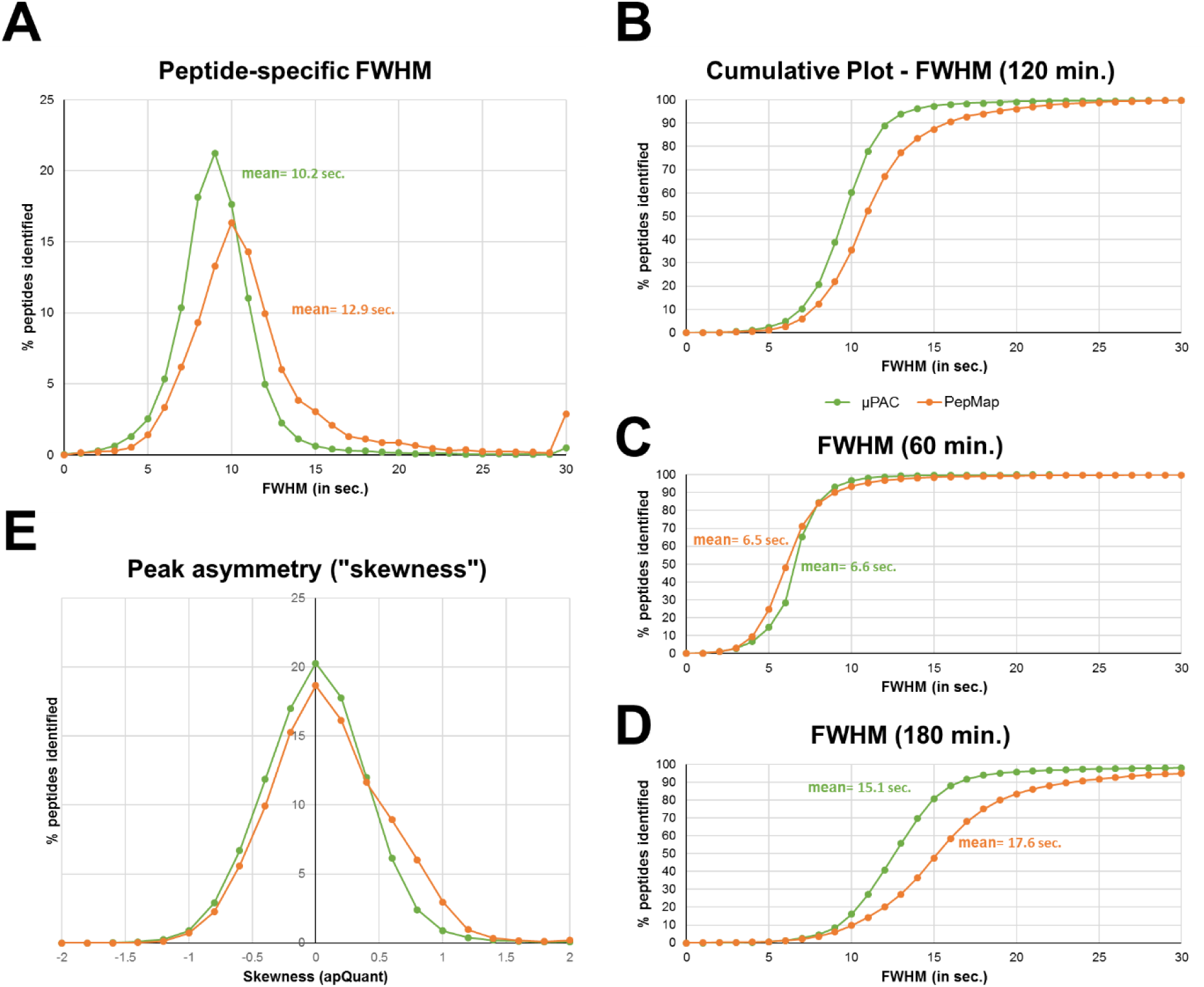
Chromatographic performance parameters of the µPAC RP18 and the PepMap C18 nano-HPLC systems. **A)** Relative comparison of the density distribution of peptide-specific peak widths (FWHM). **B)** Cumulative plot of peptide specific FWHM at 120min, **C)** 60 min. and **D)** 180 min. For all plots, FWHM bin width = 1 sec. **E)** Comparison of chromatographic peak asymmetry at 120 min gradient length, as calculated by “skewness” (bin width = 0.2). All chromatographic peak detection and feature calculations were performed using apQuant for Proteome Discoverer 2.3.

Most Interestingly, however, close investigation of the LC-MS/MS data generated in the course of this study revealed an unprecedented degree of retention-time stability of µPAC cartridges (Fig. 3.). For this, we calculated the deviation from the mean retention time of the identified peptide features across three technical replicates. While 95% of all features identified on the PepMap system were found to elute within a retention time window of approx. 44 sec. at 120 min. gradient length, the same proportion of features identified on the µPAC setup eluted within in a time window of approx. 4 sec. at 120 min. gradient length, and did not exceed 10 sec. at 180 min. gradient length. This outstanding retention-time stability and precision of µPAC cartridges clearly warrants future applications of this ultra-sensitive nano-HPLC setup in time-dependent LC-MS and LC-MS/MS data-acquisition regimes (e.g. scheduled tSIM or PRM workflows).

**Figure 3.**
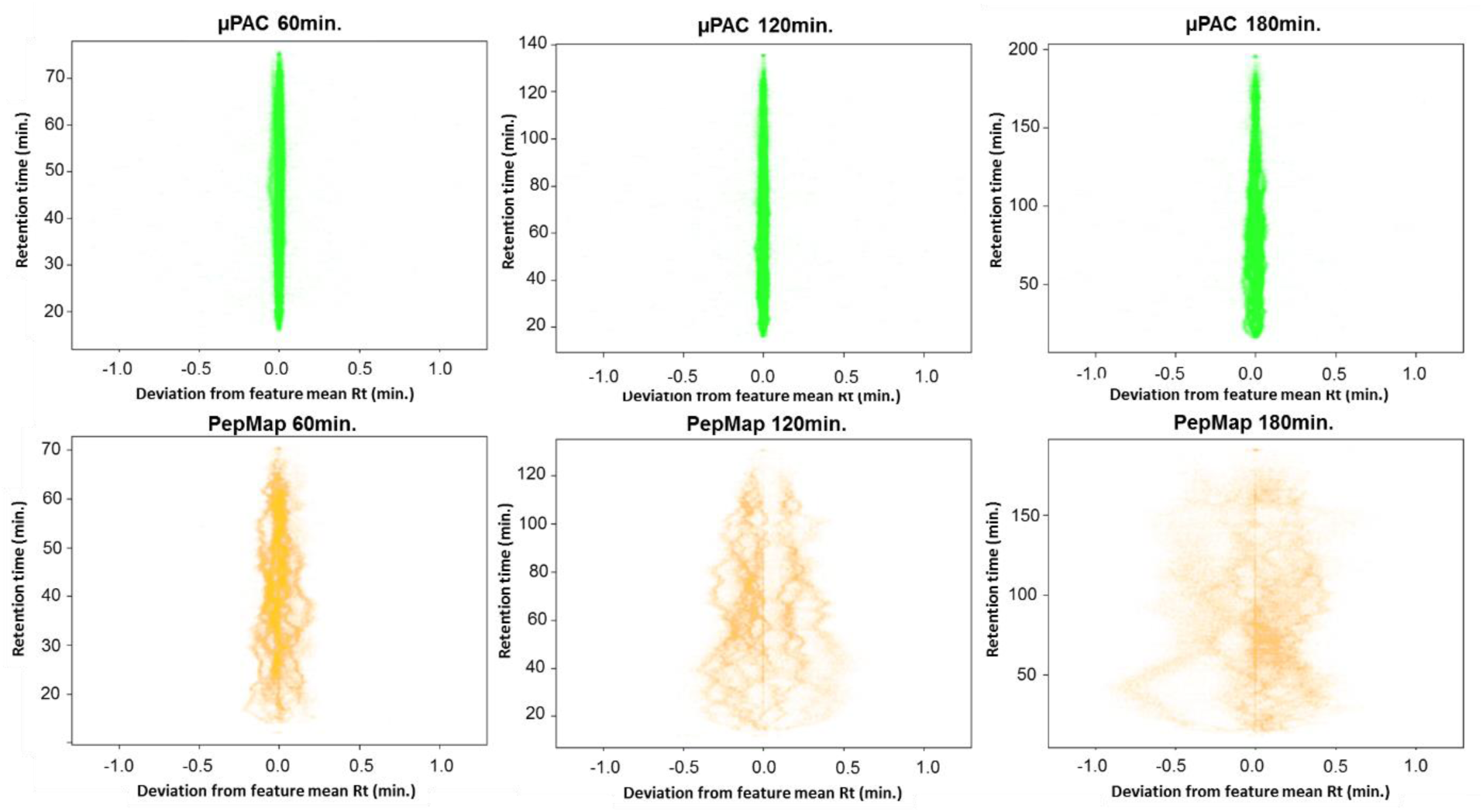
Comparison of retention time stability and precision.

Taken together, our results highlight impressive improvements in performance in the analysis of low-input proteomics samples by the application of µPAC RP18 cartridges, over current state-of-the-art nano-HPLC systems. Not only did we yield almost twice as many unique peptide and unique protein group identifications, when compared to conventional nano-flow HPLC columns, but we also observed unprecedented retention-time stability on the micropillar array-based nano-HPLC µPAC cartridges. Owing to the flexibility in design and the great potential to even further optimize chromatographic support materials and formats, this first commercial implementation of the µPAC concept clearly provides very attractive new strategies for increased sensitivity in nano-HPLC-ESI-MS/MS based proteomics platforms towards the ultra-sensitive analysis of single mammalian cells.

## SUPPORTING INFORMATION

All mass spectrometry-based proteomics data have been deposited to the ProteomeXchange Consortium via the PRIDE partner repository with the dataset identifier [PXD014124]. For review purpose please query the data using the following Reviewer account details:

Username

Password: k8kv5Isu

## ACKNOWLEDGEMENTS

We thank all members of our laboratories for helpful discussions. We acknowledge G. KrŠŠáková for method development. This work has been supported by EPIC-XS, project number 823839, funded by the Horizon 2020 program of the European Union, and the Austrian Science Fund by ERA-CAPS I 3686 International Project.

